# Neuronal aging is associated with declined autophagy caused by reduced functionality of cGAS-STING signaling

**DOI:** 10.1101/2023.09.25.559304

**Authors:** Sergio Passarella, Karina Brandes, Evelyn Dankert, Paola Cavalli, Andrea Kröger, Daniela C. Dieterich, Peter Landgraf

## Abstract

The lifelong maintenance of cognitive abilities in an increasingly aging human society is one of the major challenges of future research and medical services. For this, a better understanding of the cellular aging processes in the mammalian brain is a fundamental requirement. In particular, the functioning of postmitotic neurons, which require special strategies for lifelong functionality, is still elusive in many details. Among many other hallmarks of neuronal aging, the impairment of autophagy as an essential element of cellular homeostasis is of particular importance. However, the mechanisms for regulating these processes have not yet been fully elucidated. Establishing an *in vitro* model from primary cortical cells of the mouse brain, which shows the characteristic features of cellular senescence that are also observed in the total brain, we found the accumulation of dsDNA in the cytosol of neurons. Since dsDNA is a trigger for the activation of the cGAS-STING signaling and its primordial function is a non-canonical activation of autophagy, we analyzed its impact on aging neurons. We were able to demonstrate that the age-dependent downregulation of cGAS-STING signaling in neurons leads to an inhibition of autophagy at different levels. In contrast, activation of STING led to a complete rescue of autophagy in old neurons. Additionally, we found no evidence for age dependent cGAS-STING mediated IFN-I production. Hence, we propose that the primary function of cGAS-STING signaling in neurons is to maintain autophagy rather than contribute to age-related inflammation, and thus represents a target for therapeutic intervention.

## 1. INTRODUCTION

Long-term maintenance of cognitive independency in a worldwide continuously aging society, is one of the biggest challenges of current research. Thereby, aging itself is a vast risk factor for the occurrence of senile dementia as well as the development of Alzheimer’s (AD) or Parkinson’s disease (PD) (Herbig et al., 2006; van Deursen, 2014). It’s therefore not surprising that neuronal aging has been researched since a long time. In contrast to age-related pathologies, like neurodegenerative diseases, processes of normal brain aging are difficult to elucidate, although impairments of cerebral functions, such as cortex-and hippocampus dependent cognitive capabilities are clearly verifiable (Bettio et al., 2017; Morrison & Baxter, 2012). In particular, the fact that the brain consists of two main cell types, mitotic-active glial cells and post-mitotic, terminally differentiated neurons, implies cell-type-specific differences in aging and senescence processes. For neurons, in contrast to proliferating cell-types, this means the development of specific strategies that ensure lifelong survival even under detrimental conditions as e. g. oxidative stress or inflammation (Kole et al., 2013). Thus, the vast and complex network of processes that cause a neuron to age makes it hard to pinpoint the first step (Bishop et al., 2010). While in the most somatic cells senescence is understood as a phenomenon in which they stop dividing, this situation is already present in postmitotic neurons without expressing age related hallmarks (Kole et al., 2013). However, independently of this, neurons are able to develop a characteristic senescence-associated phenotype in the course of their aging process, which includes an increase in beta-galactosidase (SA-ß-gal) activity (Geng et al., 2010; Piechota et al., 2016), upregulation of the Cdk inhibitor p16, increasing activation of the p38 MAP kinase and loss of laminin B1 (Ishikawa & Ishikawa, 2020; Kuilman et al., 2010). Furthermore, aging neurons develop a senescence-associated secretory phenotype (SASP) (Jurk et al., 2012; Kuilman et al., 2010) that leads to the secretion of various proteins, including inflammatory cytokines. Nevertheless, the three main events of cellular aging, mitochondrial malfunction including the development of oxidative stress, DNA damage, and the decline of autophagy, also take place in neurons as well (Ishikawa & Ishikawa, 2020; Moreno-Blas et al., 2019). In addition, impairments of proteostasis, a finely balanced process consisting of protein synthesis, post-translational modification and protein degradation, also play a crucial role in the loss of neuronal function and plasticity, consequently leading to progressive neuronal aging (Ishikawa & Ishikawa, 2020). In this context, it is still elusive which of the age-dependent phenomena causes the other processes or to what extent these have cell-type-specific differences. Meanwhile, there is a general consensus that chronic, low-grade inflammation associated with SASP contributes to age-related neurodegeneration (Bussian et al., 2018; Franceschi et al., 2018; Golde & Miller, 2009; Musi et al., 2018). A growing body of evidence points to an involvement of the cyclic GMP-AMP synthase – stimulator of interferon genes (cGAS-STING) pathway leading to the generation of type-1 interferons (IFN-I) with the subsequent expression of interferon stimulated genes (ISGs) (Glück et al., 2017; Takahashi et al., 2018). The cGAS-STING axis as a part of the innate immune system elicits not only foreign DNA in order to prevent the cells from invading pathogens, but also self-DNA derived from defective mitochondria and disintegrated nuclei occurring during aging and neurodegeneration (Song et al., 2019; West et al., 2015; Zhang et al., 2014). However, the induced immune response of cGAS-STING related IFN-1 signaling does not only have cell-protecting effects. The uncontrolled stimulation of this signaling pathway leads to undesirable effects such as increased neuroinflammation along with increased neurodegeneration (Hinkle et al., 2022; Szego et al., 2022). This implies both possible endogenous control mechanisms, which still need to be further elucidated, as well as the identification of putative targets for anti-inflammatory therapy approaches (Decout et al., 2021). However, the cGAS-STING signaling pathway is not only associated with IFN-I mediated immunity but seems to be involved in several other processes (Hopfner & Hornung, 2020; Prabakaran et al., 2018; Reinert et al., 2021; Riley et al., 2018). Thus, cGAS not only catalyzes the cyclic dinucleotide GMP-AMP (2’3’-cGAMP), but also appears to have a non-catalytic function as an anti-tumor factor (T. Li et al., 2016). A whole series of functions are described for STING. In addition to the TANK-binding kinase 1 (TBK1) mediated activation of IRF3, which translocates into the nucleus after phosphorylation and dimerization and activates interferon beta 1 (IFNB1), STING leads to the activation of the nuclear factor kappa B (NF-kB) and MAPK signaling pathways (Oliveira Mann et al., 2019). In addition, STING is involved in the regulation of autophagy processes (Gui et al., 2019; Prabakaran et al., 2018; Watson et al., 2012) and the induction of lysosomal cell death and trafficking (Kuchitsu et al., 2023; Paludan et al., 2019). In particular, the STING-dependent, non-canonical induction of autophagy is of outstanding importance, since it seems to be its primordial function of eliminating microbes, and acts independently of the IFN-I response (Gui et al., 2019; Watson et al., 2012; Yamashiro et al., 2020). In the brain, the majority of STING is expressed in microglia, the resident immune cells of the brain (Paul et al., 2021). Although astrocytes and neurons also produce interferons, the role of the cGAS-STING signaling pathway in these cell types is largely unclear (Owens et al., 2014; Yamada et al., 1994).

In order to analyze the role of the cGAS-STING signaling in relation to aging processes of the brain in a cell type-specific manner, we developed an *in vitro* model system, consisting of primary neurons and astrocytes from mice. The cells were cultured for up to 60 days *in vitro* (DIV 60) and analyzed at DIV 21, DIV 40 and DIV 60. We could demonstrate that our aging culture system (ACS) represent a reliable tool for studying senescence processes cell type-specifically, seeing a tremendous pathophysiological switch in postmitotic neurons between DIV 40 and 60. Our results indicate that the age-related loss of autophagic capacity in neurons is caused by an impaired cGAS-STING signaling leading to proteostatic disbalance.

## 2. MATERIAL AND METHODS

### Animals

Male and female C57BL/6J wild type (WT) mice (Mus musculus) were kept under controlled conditions regarding temperature and humidity at the animal facilities of the Otto-von-Guericke-University Magdeburg, with a 12 h day/night cycle and with food and water ad libitum. All experiments were conducted in accordance with the recommendations of the National Committee for the Protection of Animals Used for Scientific Purposes of the Federal Republic of Germany and European Regulations for Ethical Care and Use of Laboratory Animals (2010/63/EU) and approved by the local Ethics Commission of the Federal State of Saxony-Anhalt (42502-2-1507 Uni MD and 42502-2-1578 Uni MD).

### Primary neural cultures from mice cortices and treatments

Primary cortical cells were obtained from embryonic E18-19 mice (C57BL/6J) and prepared as essentially described in Banker & Goslin (Banker & Goslin, 1988), and as published in (Müller et al., 2015). Briefly: Brains of E18.5 embryos were dissected and the cortices mechanically and enzymatically dissociated using Hank’s balanced salt solution (HBSS), containing 0,25% trypsin. After washing with HBSS, cells were diluted in Dulbecco’s modified eagle medium (DMEM) containing 10% FBS, 100 U/ml Penicillin, 100 µg/ml Streptomycin and 2 mM L-glutamine (all Gibco), and each 40.000 cells were seeded onto poly-D-lysine treated 12 mm coverslips for immunocytochemistry or each 300.000 cells per well into 6-well plates for quantitative Western-blot analysis. 3 and 24 hours later the plating medium was exchanged against NeurobasalTM containing 2% BL27 (Gibco) and 0,8 mM L-glutamine (all Gibco). The cultures were kept at 37°C, 5% CO2 and 95% humidity for 21, 40 and 60 days respectively in vitro (DIV) and fed once a week with fresh one tenth of their culture medium. To modulate the STING activity, the cells were treated for 3 h with 2’3’- cGAMP with a concentration of 12,5 µg/mL and H-151 with a concentration of 4 µg/mL. To block the fusion between autophagosome and lysosome the cells were treated for 2 or 3 h with chloroquine (CQ) with a final concentration of 50 µM.

### Immunofluorescence staining

Cells were briefly rinsed with PBS, pH 7,4, containing 1 mM MgCl2 and 0,1 mM CaCl2 (PBS-MC), and subsequently incubated with Periodate-lysine-paraformaldehyde (PLP) fixative for 30 min at RT (according to (McLean & Nakane, 1974). Following three times washing with PBS, pH 7,4, cells were blocked in B-block solution (10% normal horse serum, 5% sucrose, 2% BSA, 0,2% Triton X100 in PBS, pH 7,4) for one hour at RT. Afterwards the cells were incubated with the respective primary antibodies diluted in B-block solution overnight at 4°C. Next, cells were washed three times with PBS, pH 7,4, at RT and again incubated with B-block solution containing the respective fluorophore-conjugated secondary antibodies for one hour at RT. Finally, cells were washed three more times with PBS, briefly rinsed with ddH2O and subsequently mounted with Mowiol (Calbiochem). If a nuclear staining was required, the least washing step was replaced by 10 min incubation with 1:5000 DAPI in PBS, pH 7,4. Cells were imaged using a confocal laser scanning fluorescence microscope (CLSM; Axio observer.Z1 microscope provided with a LSM 710 and LSM 800 confocal system; Carl Zeiss group). All images were taken as 8-bit Z-stacks with 0.08 μm pixel size using a Plan-Apochromat 63x/numerical aperture 1.4 oil DIC M27 immersion objective, except for the Sholl analysis were an EC Plan Neofluar 20x was used. The confocal microscope images were processed as maximum intensity projections confocal pictures (89,97×89,97 µm – 1024×1024 pixels) and analyzed in FIJI (version 2.0). For intensity analysis, the intensity values of each cell were related to the respective mean intensity. The Sholl analysis of neurons was carried out using the plugin Simple Neurite Tracer (SNT) created by Prof. Tiago A. Ferreira (Arshadi et al., 2021) included in the FIJI software.

The following primary antibodies were used:

cleaved caspase 3 (Cat. No. MAB835, 1:200 in blocking solution, R&D Systems), dsDNA (Cat No. ab27156, 1:300 in blocking solution, Abcam, Cambridge, MA), GM130 (Cat. No. 610822, dilution 1:500 in blocking solution, BD Bioscience), LAMP1 (Cat No. ab208943, dilution 1:300 in blocking solution, Abcam, Cambridge, MA), MAP2 (Cat No. 188004, dilution 1:1000 in blocking solution, Synaptic Systems), p62 (Cat. No. ab56416, dilution 1:300 in blocking solution, Abcam, Cambridge, MA), STING (Cat. No. 19851-1-AP, dilution 1:300 in blocking solution, Proteintech).

### Metabolic labeling and fluorescent non-canonical amino acid tagging (FUNCAT)

For metabolic labeling of nascent proteins, the non-canonical methionine (Met) surrogate azidohomoalanine (AHA) was used in a final concentration of 4 mM. Therefore, the culture medium was replaced by Met-free Hibernate medium and after 30 min pre-incubation time supplemented with AHA. As a control, cells were supplemented with an equimolar Met concentration. After 3 hours cells were briefly rinsed with PBS-MC and finally fixed using 4% PFA. The FUNCAT reaction was performed as described previously (Dieterich et al., 2010; Müller et al., 2015). In brief: Cells were washed three times with PBS, pH 7,4, subsequently blocked at RT with B-blocking solution for 1,5 h under gently agitation and washed again three times using PBS, pH 7,8. Next, the click reaction mix was prepared in PBS, pH 7,8, containing 20 µM Triazole ligand (Tris[(1-benzyl-1H-1,2,3-triazol-4- yl)methyl]amine; TBTA, Sigma-Aldrich), 500 µM TCEP (Tris(2-carboxyethyl)phosphine hydrochloride; Sigma-Aldrich), 0,2 µM 5/6 TAMRA-PEG4-Alkyne tag (Jena Biosciences) and 0,2 mM freshly prepared CuSO4. Note, the reaction mix must be sequentially vortexed for 10 sec after adding each component. Finally, each 400 μl/well of the reaction mix were transferred to the fixed cells and incubated in completely darkness over night at RT. The click reaction was stopped via three times washing using 0,5 mM EDTA and 1% Tween-20 in PBS, pH7,4, followed by three times washing with PBS, pH7,4. Afterwards, the cells can be further treated for immunofluorescence counterstaining.

### Lysotracker staining

For the staining of lysosomes in living cells Lysotracker Red DND-99 in a final concentration of 50 nM (Thermo Fisher cat. no. L7528) was used according to the manufacturer’s instructions.

### 2’,3’-cGAMP measurement

For measuring 2’3’-cGAMP, a competitive ELISA kit (Item No 501700, Cayman Chemical) was used following the manufacturer’s instructions and analyzed at a wavelength of 450 nm using an Infinite F50 (Tecan) photometer.

### Terminal deoxynucleotidyl transferase dUTP nick end labeling (TUNEL) assay

The TUNEL assay was caried out using the In Situ Cell Death Detection TMR rot kit (Roche, cat. no. 12156792910) according to the manufacturers protocol. As a positive control, cells were treated with DNase I for 10 min at 37°C in Order to induce DNA strand breaks.

### Senescence ß-galactosidase cell staining

ß-galactosidase levels were analyzed using the senescence ß-galactosidase staining kit (Cell Signaling) according to the manufacturer’s specification (Cell Signaling, cat. no. #9860). Cells were briefly washed with ice-cold PBS, pH 7,4, and subsequently incubated for 15 min in the fixative solution of the kit. Following three times washing with PBS, pH 7,4, cells were incubated at 37°C overnight in the ß-Galactosidase staining solution. Terminal, cells were washed three times with PBS, pH 7,4, rinsed with ddH2O and subsequently mounted with Mowiol. Images were obtained using a Leica DMS1000.

### Protein extraction

For sodium dodecyl sulphate-polyacrylamide gel electrophoresis (SDS-PAGE) and Western-blot analysis primary cortical cells were grown in 6 well plates at a density of 300.000 cells/well. At indicated timepoints, cells were washed with 1 ml PBS-MC and subsequently scraped and directly lysed with protein sample buffer (62,5 mM Tris-HCl, pH 6.8, 1% SDS, 10% glycerol, 5% ß- mercaptoethanol, 0,001% bromophenol blue). The lysates were finally incubated at 95°C for 5 min, cooled on crushed ice and stored at -20°C. The respective protein concentrations were determined using the amido black assay according to Badin & Herve (BADIN & HERVE, 1965).

### SDS-PAGE and Western blot

SDS-PAGE was conducted according to Laemmli (Laemmli, 1970) using 5-20% gradient gels in a Hoefer™ Mighty Small System (Fisher Scientific). For each sample a total of 10 µg protein was loaded and separated at a voltage of 12 mA per gel and a constant temperature of 4°C. If required, gels were stained using Coomassie Brilliant Blue R250. Otherwise, proteins were transferred to nitrocellulose (NC) membranes (Protran BA85, 0,22 µm, Li-Cor Biosciences) using the Hoefer™ Mighty Small System SE250 (Fisher Scientific) at 200 mA for 90 min at a constant temperature of 4°C. Subsequently, the membranes were stained with Ponceau staining solution (0,5% Ponceau S, 3% acetic acid in dd H2O) for 10 min and after destaining the membranes were incubated at 4°C over night with primary antibodies in the respective concentration using 1xTBS containing 0.1% Tween 20 including LC3 A/B (Cat. No. #4108S, dilution 1:1000 in blocking solution, Cell Signaling Technology), p62 (Cat. No. #5114, dilution 1:1000 in blocking solution, Cell Signaling Technology), GFAP (Cat. No. 170002, dilution 1:1000 in blocking solution, Synaptic Systems), MAP2 (Cat. No. M4403, dilution 1:1000 in blocking solution, Sigma-Aldrich), TAU1 (Cat. No. MAB3420, dilution 1:1000 in blocking solution, Sigma-Aldrich), GST (Cat. No. ab53942, dilution 1:1000 in blocking solution, Abcam, Cambridge, MA), TOM20 (Cat. No. #42406, dilution 1:1000 in blocking solution, Cell Signaling Technology), cGAS (Cat. No. #31659, dilution 1:1000 in blocking solution, Cell Signaling Technology), STING (Cat. No. 19851-1-AP, dilution 1:300 in blocking solution, Proteintech), TBK1 (Cat. No. #29047, dilution 1:1000 in blocking solution, Cell Signaling Technology), pTBK1 (Ser172) (Cat. No. #5483, dilution 1:1000 in blocking solution, Cell Signaling Technology), IRF3 (Cat. No. ab68481, dilution 1:1000 in blocking solution, Abcam, Cambridge, MA), pIRF3 (Ser396) (Cat. No. PA5-38285, dilution 1:1000 in blocking solution, Invitrogen). After washing the blots were incubated for 90 minutes at room temperature with HRP-conjugated secondary antibodies (1:7500) and finally imaged in a LICOR OdysseyFC (LI-COR). Protein quantities were analyzed using Image Studio Lite software (LI-COR® Biosciences GmbH, version 5.2.5). Normalization was carried out on the control group and related to the corresponding ß-actin (monoclonal mouse, Cat. No. #3700, dilution 1:2000 in blocking solution, Cell Signaling Technology and monoclonal rabbit, Cat No. #8457, dilution 1:2000 in blocking solution, Cell Signaling Technology) values.

### Quantification and Statistical Analysis

Data are shown as mean ± SEM. One-way analysis of variance (ANOVA) followed by post hoc Tukey’s multiple comparison test was used to determine statistical differences among three or more experimental groups for Western blot and images data. Two-way ANOVA followed by Holm–Šídák test was conducted for Sholl analysis of cortical neurons. Differences between mean values P < 0.05 were considered statistically significant. All present data were analyzed and graphed with Prism 8 (GraphPad Software Inc. La Jolla, CA, USA).

## 3. RESULTS

### Primary cortical neurons from mice develop a senescence phenotype after DIV 21 followed by apoptotic features after DIV 40

Primary neural cultures are obtained from the cortices of mouse embryos at the developmental stage E18.5. Already after few days, the co-cultures consist predominantly of neurons and astroglia and are mostly devoid of microglial cells. For our experiments we used these in vitro cultures as a model system for the cell type specific analysis of aging processes and apoptosis in neurons. Up to 40 days in vitro, we could observe no or only minor changes in the cellular composition of our model system (Figure S1 a-d). In contrast, between DIV 40 and 60 the GFAP concentration increased significantly, indicating increased proliferation of astrocytes in the culture, while in the same time period a significant decrease in the neuronal markers MAP2 and TAU1 was observed, indicating both a morphological alteration of the neurons or their partial loss. Sholl analysis revealed morphological changes in neurons, such as a decrease in dendrite length and branching, starting between DIV 40 and 60 (Figure 1 a-c) and are consistent with previous reports (Burke & Barnes, 2006; Ishikawa & Ishikawa, 2020). As one of the earliest signs of cellular stress and the development of a senescence phenotype in neurons, we observed a significant increase in ß-gal positive cells between DIV 21 and 40 (Figure 1 d, Figure S2 a). However, the development of the senescence phenotype between DIV 21 and 40 was characterized by only moderate but significant cell loss through apoptosis as detected via TUNEL assay (Figure 1 e, Figure S2 b), mainly due to the fact that only negligible amounts of dead cells were found for DIV 21. In contrast, the number of apoptotic cells was drastically increased at DIV 60, indicating a switch from senescence to apoptotic processes between DIV 40 and 60 (Figure 1 e, Figure S2 b). Other markers for cellular stress and senescence were also only slightly increased up to DIV 40 and augmented significantly between DIV 40 and 60. For example, the expression of glutathione S transferase (GST), as a characteristic feature of high reactive oxygen species (ROS) concentrations (Raza, 2011), was only slightly increased up to DIV 40, but almost doubled up to DIV 60 (Figure S1 a, e), while the number of mitochondria as the main source of ROS production became first significant decreased up to DIV 60 (Figure S1 a, f). Furthermore, we observed the accumulation of poly-ubiquitin-conjugated proteins as a hallmark of the senescence-related decreasing ability of protein degradation and recycling (Kirkin et al., 2009). In our cultures ubiquitination showed a tendency to increase already at DIV 40, with a significant boost up to DIV 60 (Figure 1 f; Figure S2 c). The observed mitochondrial stress, which ultimately leads to mitochondrial disintegration and loss of the organelles, is associated with increasing accumulation of double-stranded DNA (dsDNA) in the cytosol of neurons, beginning before DIV 40 and peaking at DIV 60 (Figure 1 g, h, i). However, in this context we cannot clearly state that the accumulation of dsDNA in the cytosol that we observed consists exclusively of mitochondrial DNA (mtDNA), but also contains components of nuclear origin. Irrespective of this, we see a direct correlation between the observed age-dependent degradation of mitochondria and the continuous increase in activated caspase 3 in the cytosol of the cortical primary neurons (Figure 1 j, k), most probably caused by the release of cytochrome C from the mitochondria (Kluck et al., 1997), which reaches its maximum at DIV 60 and points towards a switch from the senescence phenotype towards an apoptotic phenotype between DIV 40 and 60.

**FIGURE 1.**
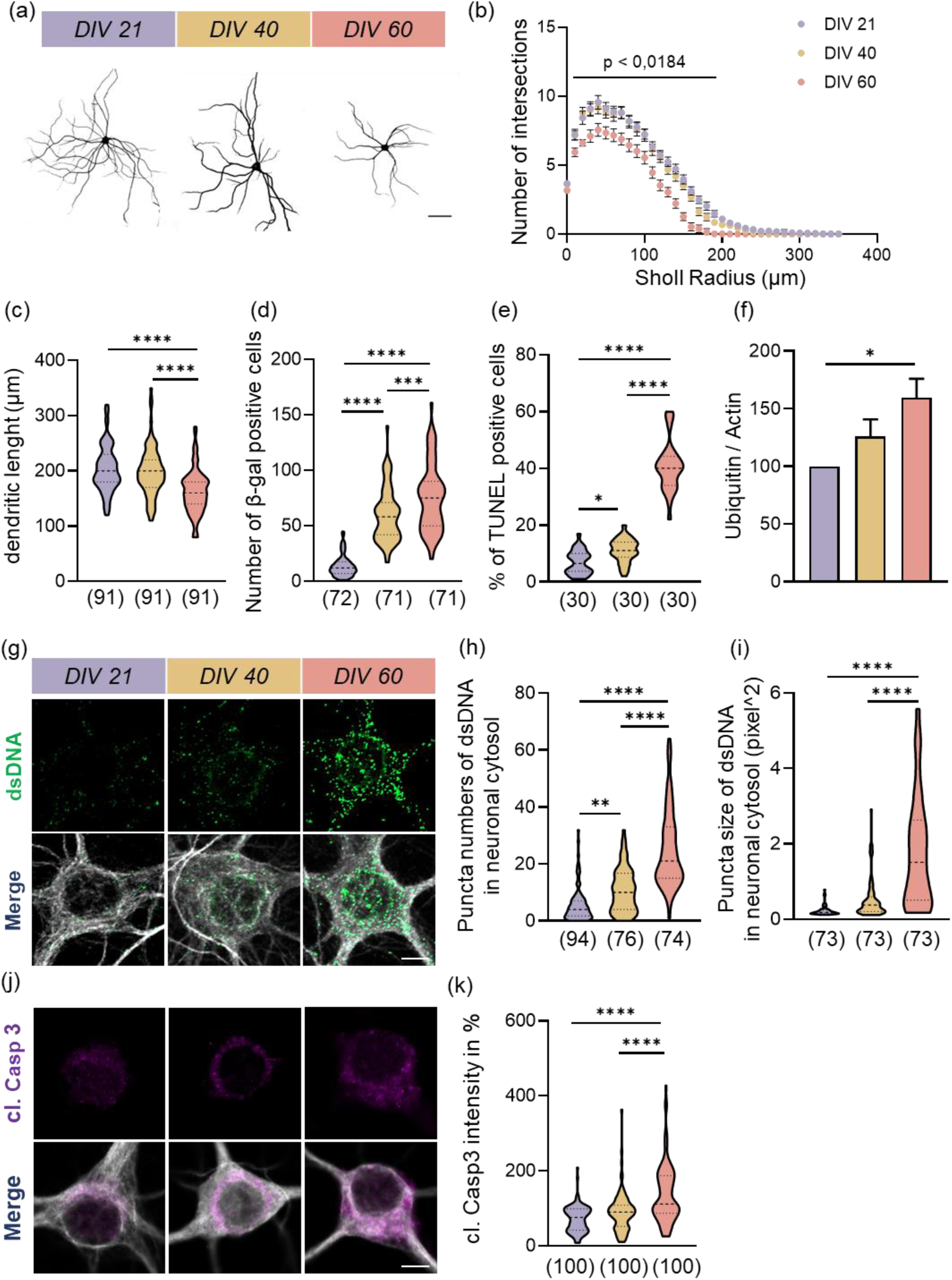
Primary E 18.5 mouse neurons at DIV 60 display features associated with physiological brain aging including morphologic structural changes in comparison with younger cultures. (a) Neurons were stained with MAP2 (black shape) in order to evaluate morphological changes during different time points (DIV 21, DIV 40 and DIV 60), scale bar = 20 µm. (b, c) Quantification of Sholl analysis shows a significant reduction of dendritic arborisation and length in aged neurons compared with neurons at DIV 21 and DIV 40. Individual values obtained from n = 22-23 neurons/group from four independent experiments. (d) Beta-galactosidase activity was analysed at DIV 21, 40 and 60. Each data in the graph represent the number of cells positive beta-galactosidase staining for single picture (size: 2195 µm x 1737 µm) (representative pictures shown in supplementary data 2). Graph represents n = 17-18 images/group from four independent experiments. (e) Relative number of TUNEL positive cells evaluated on the total amount of DAPI signals. Graph shows n = 10 images/group from three independent experiments. The size of TUNEL assay pictures is 850,19 x 850,19 µm (representative pictures are depicted in supplementary data 2). (f) Quantitative data of immunoblotting analysis of ubiquitinated proteins during aging *in vitro* normalized to Actin. Values are the mean of three technical replicates in each group from three independent experiments for immunoblotting analysis. (g) Immunostaining of dsDNA puncta (green channel) on somàs neurons labelled with MAP2 (white channel) in DIV 21, 40 and 60 cortical culture, scale bar = 5 µm. (h, i) Graphs show data analysis evaluating the number and size (evaluated in pixel^2^) of dsDNA staining. Graphs show n = 15-24 images/group from three or four independent experiments. (j, k) Representative pictures of cleaved caspase 3 (magenta channel) immunofluorescence staining, and quantification of fluorescence intensity of cleaved caspase 3 positive neurons stained with MAP2 (white channel), scale bar = 5 µm. All data are presented as the mean ± SEM. Symbols for P-values used in the figures: \**P* < 0.05, \*\**P* < 0.01, \*\*\**P* < 0.001, \*\*\*\**P* < 0.0001. P-values were determined by ordinary one-way ANOVA with Tukey’s multiple comparisons test and for Sholl analysis data were analyzed with two-way ANOVA with Tukey’s multiple comparisons test.

### Protein synthesis capacity is noticeably impaired in senescent primary cortical neurons

Since cellular proteostasis is closely linked to aging processes, we examined next the protein synthesis capacity of aging neurons in a primary cortical culture. For this purpose, we used the fluorescence non-canonical amino acid tagging (FUNCAT) technology, which was developed and constantly refined in our laboratory (Dieterich et al., 2010). Already after three hours of metabolic pulse labeling with AHA and subsequent click-chemistry using an alkyne-carrying TAMRA fluorophore, we obtained a rich fluorescence signal both in the soma and in the proximal main dendrites of 21-day-old, MAP2-positive neurons, which indicates an adequate protein synthesis rate (Figure 2 a, b). A comparison of the protein synthesis rates of DIV 21, -40 and -60-day-old primary neurons revealed that the rate of newly synthesized proteins in the soma of the cells was already significantly reduced at DIV 40 and more than 50% less at DIV 60 (Figure 2 a, c). Also, in the area of the proximal main dendrites, the amount of newly synthesized proteins was already significantly reduced at DIV40 and continued to decrease up to DIV60 (Figure 2 b, d). This leads to the conclusion that in neurons of primary cortical cultures, the protein synthesis capacity decreases age-dependent dramatically between DIV21 and DIV60. At this point it is not possible to state whether the ability for local protein synthesis in the dendrites is affected in the same way. But regardless, the observed age-dependently impaired protein synthesis capacity, together with the reduced ability of proteasome-and autophagy-dependent protein degradation, fits into the overall picture of a senescence related increasing defective proteostasis in our in vitro model of cortical primary cultures and makes it a useful subject for our following studies.

**FIGURE 2.**
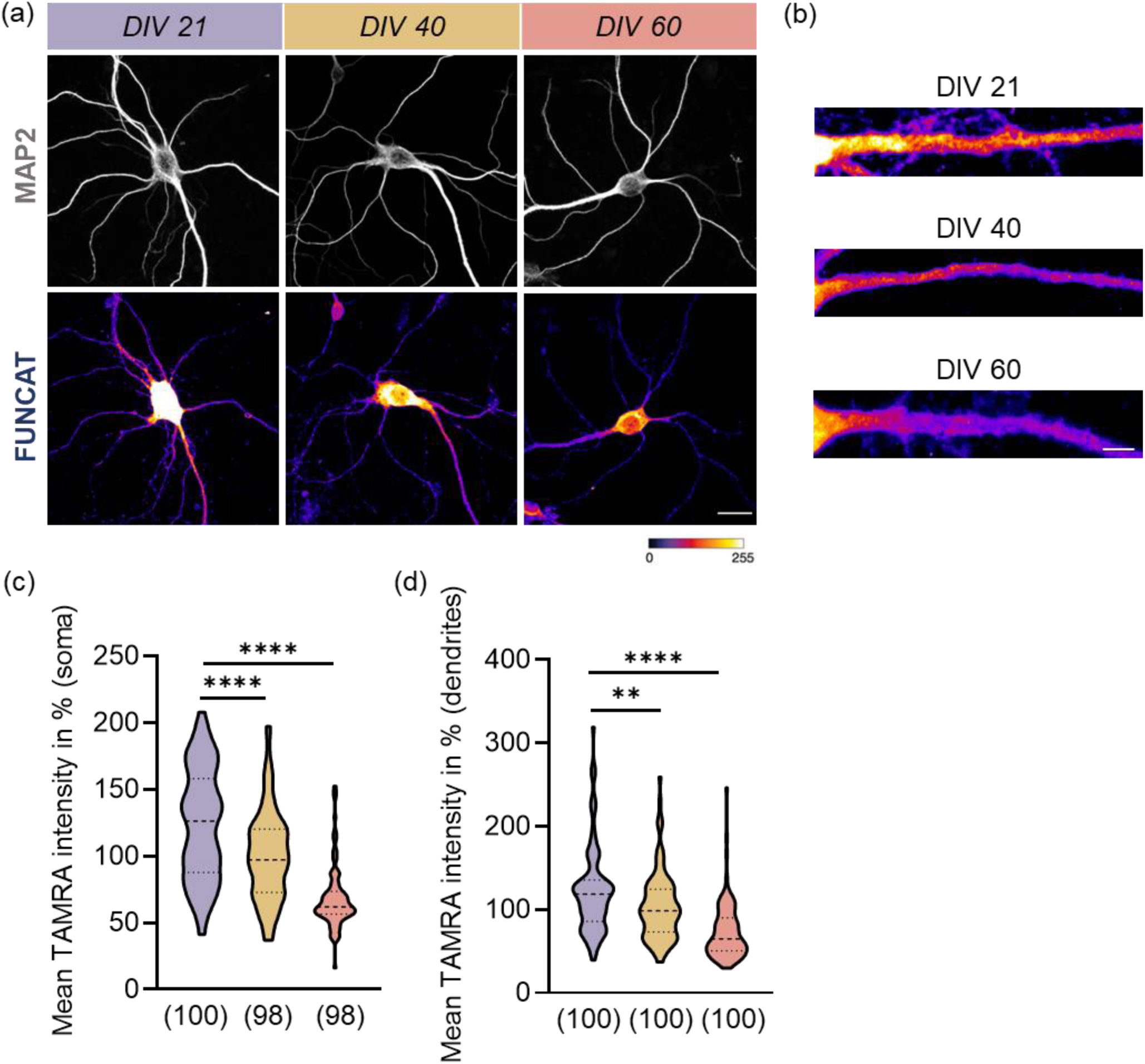
Decrease of the *de novo* protein synthesis in old neurons. Neurons were metabolically labelled with the non-canonical amino acid AHA 4 mM for 2 h in Hibernate at different time point DIV 21, 40 and 60. (a) Immunostainings of MAP2 (white channel) with FUNCAT signal. (b) Enlarged image to appreciate the changes in the TAMRA signal intensity at the dendritic level. (c,d) Quantification of TAMRA fluorescence intensity on soma and on proximal dendritic segments. All data are presented as the mean ± SEM, 24 - 25 figures done for each group from four independent experiments for FUNCAT-analysis. Symbols for P-values used in the figures: \*\**P* < 0.01, \*\*\*\**P* < 0.0001, scale bar = 20 µm (a), scale bar = 1 µm (b). P-values were determined by ordinary one-way ANOVA with Tukey’s multiple comparisons.

### Autophagy and lysosomal proteolysis pathway are impaired in senescent and apoptotic primary cortical cultures

Age-related impairments in cellular autophagy have far-reaching consequences for proteostasis in primary neuronal cultures (Ishikawa & Ishikawa, 2020; Moreno-Blas et al., 2019). Therefore, we first examined the most commonly used markers for autophagy processes, LC3 and p62. While the inactive form of LC3 (LC3-I) remained constant over a period of DIV 60 (Figure S3 a, b), we observed in the same time frame a continuous increase in the phosphatidylethanolamine-conjugated and thus membrane-integrated form of LC3 (LC3-II) (Figure S3 a, c). The resulting, age-dependent, increased ratio of LC3-II and LC3-I (LC3-II/LC3-I) at DIV40 and 60 indicates an early impairment of autophagy in primary cortical cultures (Figure S3 d). In contrast, we found no accumulation of the ubiquitin-binding autophagic adapter protein p62 at DIV40 and only a slight accumulation on DIV 60 (Figure S3 a, e). To analyze p62 cell-specific in neurons, we performed immunofluorescence staining and quantified the p62-positive signals within the soma of a MAP2 mask (Figure 3 a, c). Thereby, we were able to show that p62 accumulation in the primary cortical neurons does not occur until DIV 60, which indicates a relatively late impairment of autophagy in this specific cell type (Figure 3 b). More detailed analysis of all experimental age groups using CQ, an autophagosome-lysosome fusion blocker (Mauthe et al., 2018), revealed that equal amounts of p62-positive autophagosomes accumulated in the soma of neurons, regardless of the respective neuron ages (Figure 3 c, d). This finding indicates a senescence-related reduction in phagosome-lysosome fusion rather than altered autophagosome biogenesis. Since there is a direct functional relationship between autophagy and the lysosomal proteolysis pathway, we analyzed the number of lysosomes in neurons using the marker protein LAMP1. Starting at DIV 40, we observed a continuous decrease in LAMP1-positive puncta that reached its minimum at DIV 60, indicating a senescence-dependent reduction in lysosome biogenesis (Figure 3 e, f). Surprisingly, we found significantly enlarged LAMP1 puncta for neurons at DIV 60, implying an age-dependent increase in lysosome size (Figure 3 g). In order to gain more information about the functionality of the observed neuronal lysosomes, we used a fluorescence-labeled lysotracker, which only detects functional lysosomes with an acidic environment (Figure 3 h). We found an age-dependent significant increase in lysotracker-positive signals with a maximum at DIV 60 (Figure 3 i), pointing towards a higher lysosomal capacity in the older neurons. Regardless of the fact that further investigations into the reasons leading to an age-dependent increase in lysosome formation with a simultaneous decrease in LAMP1 are required, the prevention of autophagosome-lysosome fusion and degradation seems to be responsible for the impaired autophagy-dependent protein degradation.

**FIGURE 3.**
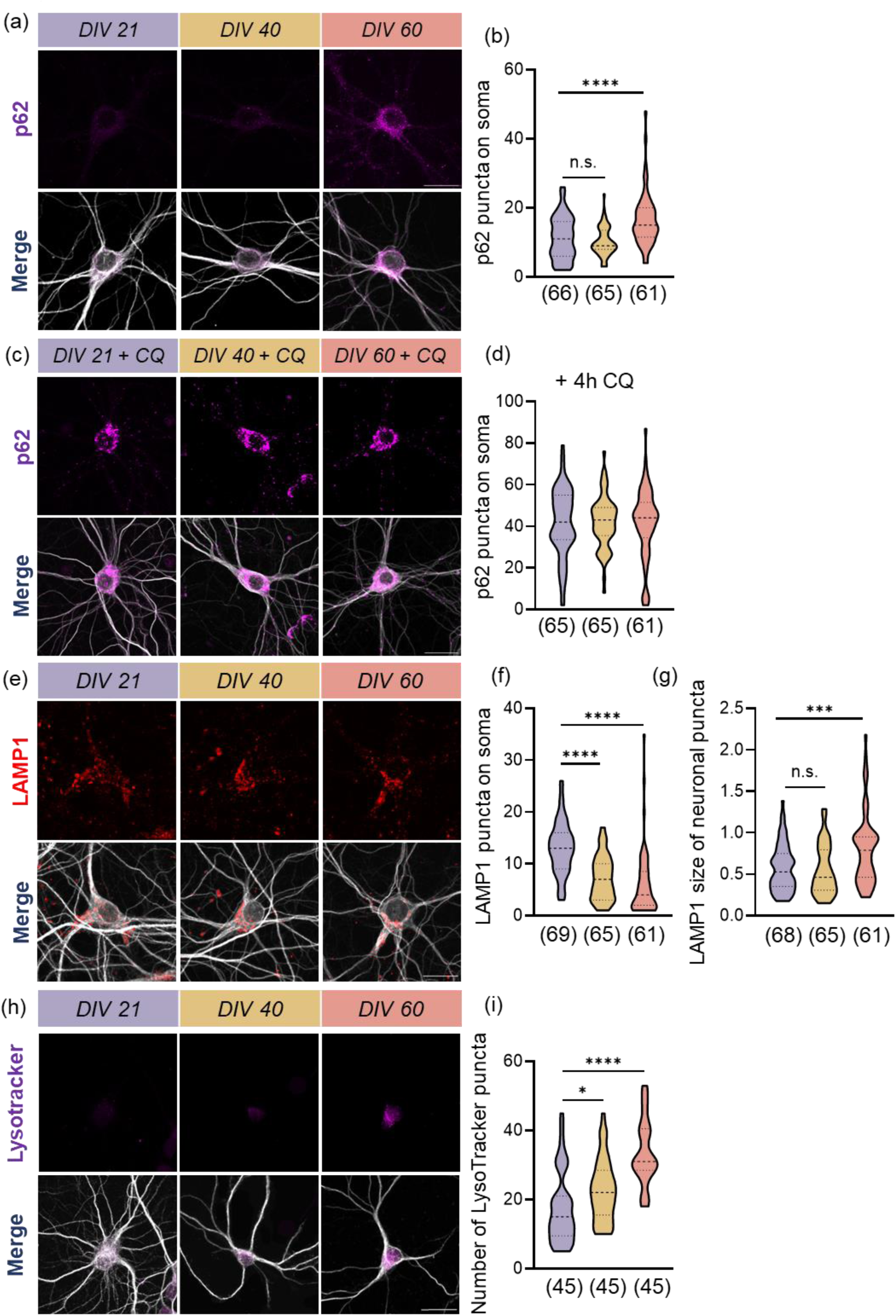
Autophagic impairment and dysfunctional lysosomal homeostasis contribute to neuronal aging *in vitro*. At DIV 60 neurons show an accumulation of p62 positive autophagosomes and neclectible accumulation of autophagosomes after CQ treatment. (b,d) Neurons were stained with p62 (magenta) and MAP2 (white) at different time point DIV 21, 40 and 60, without (a) and with CQ (c) and assessment of the numbers of p62 puncta, scale bar = 20 µm. (b,d) Analysis of p62 puncta on neuronal soma without and with the CQ treatment in order to define the autophagic situation. Graphs show n = 20-22 images/group from three independent experiments. (f) Cortical mouse neurons with different age, were immunostained with LAMP1 (red channel) and MAP2 (white channel), scale bar = 20 µm. (f, g) Quantitative analysis of number and size (in pixel^2^) of LAMP1 puncta on the neuronal soma. Individual values obtained from n = 20-23 neurons/group from three independent experiments. (h) In order to evaluate the number of active lysosomes, cortical neurons were labelled with LysoTracker Red DND-99 (magenta) and MAP2 (white) during aging *in vitro*. (i) Evaluation of LysoTracker puncta on neuronal soma with n = 15 images/group from three independent experiments. All data are presented as the mean ± SEM. Symbols for P-values used in the figures: \*\*\**P* < 0.001, \*\*\*\**P* < 0.0001. P-values were determined by ordinary one-way ANOVA with Tukey’s multiple comparisons.

### The cGAS-STING pathway is downregulated in senescent primary cortical cultures

The cGAS-STING signal transduction pathway is an essential and meanwhile intensively studied component of the innate immune system (Burdette et al., 2011; X.-D. Li et al., 2013; Sun et al., 2013). However, while the cGAS-STING signaling axis is relatively well understood within the immune system and peripheral tissues, many of its functions in the CNS remain to be unraveled (Paul et al., 2021). In particular, the fact that its uncontrolled activation would lead to a permanently increased level of interferon type 1 and thus to elevated neuroinflammation and -degeneration, makes a reliable control of the cGAS-STING signaling indispensable (Hinkle et al., 2022; Sharma et al., 2020, Gulen et al. 2023). In the brain, STING is mainly expressed in microglia, the resident immune cells of the brain (Reinert et al., 2016, Gulen et al. 2023), and thus most findings relate to this cell type. Therefore, our in vitro model system from primary co-cultures of the cortex is particularly suitable for investigating the functions of the cGAS-STING signaling pathway during aging processes in neurons, as it does not contain any microglia. First quantitative analyzes of the key proteins cGAS and STING in our primary cortical cultures revealed age-dependent decreasing level of cGAS, which were only 25% of the initial amount at DIV 60, while the level of STING remained unchanged during neuronal aging (Figure 4 a-d). Since the cell-specific expression of STING in neurons are rather contradictory (Fritsch et al., 2023), we performed immunofluorescence staining with primary cortical cultures generated from wild-type and STING knockout mice and showed that STING is indeed expressed in neurons (Figure 4 k). To investigate the consequences of the age-dependent decrease of the cGAS levels, we analyzed next the levels of 2,3-cyclic GMP-AMP (2’3’-cGAMP) in the supernatant of the primary cultures using ELISA. Consistent with the age-dependent decrease in the amount of the enzyme cGAS, we also found decreasing amounts of 2’3’- cGAMP in the supernatant of the primary cultures (Figure 4 i). Blocking of the 2’3’-cGAMP receptor STING using the specific inhibitor H-151 led to a drastic accumulation of 2’3’-cGAMP in the supernatant of DIV21 cultures, indicating a prevented binding to STING and the associated recycling (Figure 4 j). Since supplementation of DIV40 old primary cultures with the STING antagonist H-151 led to comparable high levels of 2’3’-cGAMP in the supernatant, it can be assumed that cGAS has still unrestricted enzyme activity at this timepoint. In contrast, the significantly reduced 2’3’-cGAMP accumulation in the supernatant of DIV60 old primary cultures supplemented with H-151 indicates an age-dependent, already clearly restricted cGAS function (Figure 4 j). Since STING activation by 2’3’-cGAMP, as its best investigated function, leads to phosphorylation of interferon response factor 3 (IRF3), we examined the expression and activation of TANK binding kinase 1 (TBK1), which in turn phosphorylates IRF3 after binding the activated STING, and IRF3. For TBK1, we found an age-dependent trend towards a decrease in both the concentration of the total protein and its phosphorylated form (Figure 4 b, e, f). In contrast, we found slightly but not significantly increased concentrations of IRF3 in older cultures, while the phosphorylated and thus activated form of IRF3 already halved at DIV40 and was only about 25% at DIV60 (Figure 4 b, g, h). The age-dependent, drastically reduced IRF3 activation thus correlates with the amount of cGAS, which has been reduced by comparable values as described at the beginning, and the associated lower 2’3’-cGAMP concentrations. This suggests that senescence-related down-regulation of IRF3 phosphorylation via the cGAS-STING pathway in aging primary cortex cultures effectively suppresses increased IFN-I production and a consequently associated inflammatory phenotype.

**FIGURE 4.**
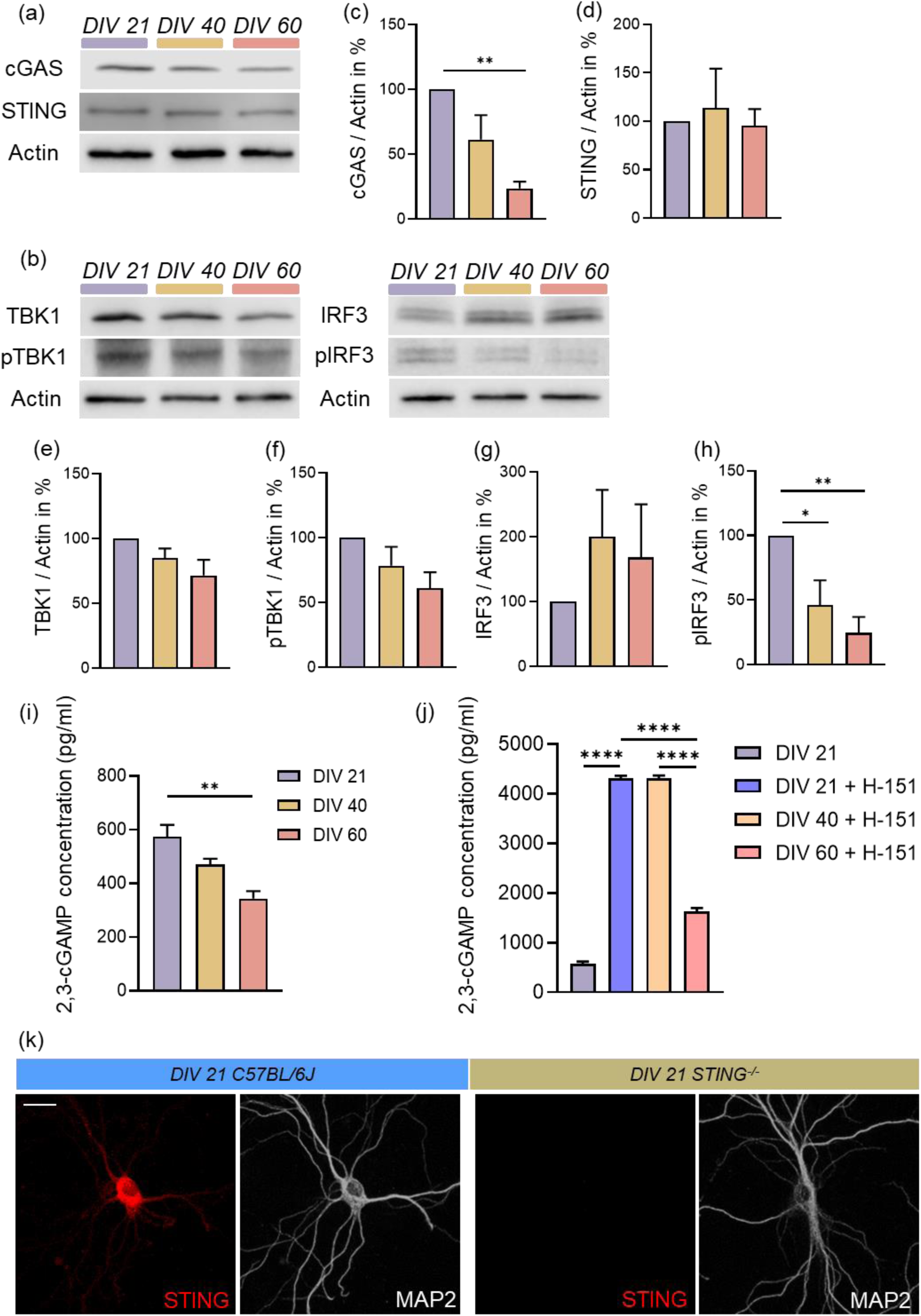
Reduced protein levels and activity of the principal protagonist of the cGAS-STING pathway during aging *in vitro* and demonstration of the presence of STING in neurons. (a, b) Representative blots of proteins involved in cGAS-STING signaling and Actin as an internal control. (c-h) Determination of selected protein levels related to the pathway using Western Blots. Relative quantities were normalized to Actin. Values are the mean of three technical replicates in each group from four independent experiments for immunoblotting analysis. (i) 2’3’-cGAMP level were measured by ELISA in the medium of neural cultures at different ages. (j) Cortical neurons at DIV 21, 40 and 60 were treated with H-151 (4 µg/mL) for 3 h, and the 2’3’-cGAMP (12,5 µg/mL) was evaluated using ELISA. 2’3’-cGAMP ELISA data were obtained from 3 independent primary neural cultures. (k) Representative pictures of WT and STING^-/-^ cortical neurons at DIV 21 stained with STING (red channel) and MAP2 (white channel), scale bar = 20 µm. All data are presented as the mean ± SEM. Symbols for P-values used in the figures: \**P* < 0.05, \*\**P* < 0.01, \*\*\**P* < 0.001, \*\*\*\**P* < 0.0001. P-values were determined by ordinary one-way ANOVA with Tukey’s multiple comparisons.

### Impaired STING mediated signaling contributes to dsDNA accumulation and proteostasis failure in senescent primary cortical neurons

Probably the best-studied function of STING is its capacity to trigger IRF3 phosphorylation and thus contribute to the expression of numerous antiviral genes via increased production of IFN-I and other proinflammatory cytokines. However, a number of other, evolutionarily more ancient functions of STING have also been described, including NF-kB activation, regulation of autophagy and the induction of lysosomal cell death (Gui et al., 2019); for review see (Hopfner & Hornung, 2020). Since we observed the accumulation of dsDNA in the soma of neurons (Figure 1g-i) with simultaneous impairment of autophagy and lysosomal processes (Figure 3 a-i) in our aging primary cortical cultures, but on the other hand a corresponding decrease in 2’3’-cGAMP and pIRF3, we analyzed next to what extent cGAS-STING signaling is involved in these processes. In the first set of experiments, we supplemented our cultures at the respective ages for three hours with the STING agonist 2’3’-cGAMP and analyzed the level of dsDNA in the soma of neurons. While in the group of the youngest neurons (DIV21) we found no change in dsDNA levels after 2’3’-cGAMP treatment, we observed a significant decrease in dsDNA levels in the DIV40 and DIV60 old neurons after 2’3’- cGAMP application (Figure 5 a, b, Figure S4 a, b). In contrast, blocking the STING activity via its specific inhibitor H-151 leads to a strong, significant accumulation of dsDNA in the area of the neuronal soma already in DIV21 old primary cultures, but also in DIV40 and DIV60 old ones (Figure 5 a, b, Figure S4 a, b). From this observation it can be concluded that STING is directly involved in the removal of dsDNA from the neuronal soma and its reduced functionality during aging is most likely linked to the age-dependent degradation of cGAS and the consequent decreasing level of 2’3’- cGAMP. Furthermore, we analyzed the impact of STING on the autophagic flux of aging neurons by monitoring the levels of the selective autophagy receptor p62 in the soma of neurons. Activation of STING using 2’3’-cGAMP already led to a slight but significant decrease of p62 in the soma of neurons from DIV21-old primary cultures, which indicates an improvement in autophagic flux (Figure 5 d; Figure S5 a). This effect was even more evident in the DIV40 and especially DIV60 old neurons, in which we observed an age-dependent strong accumulation of p62 (see also Figure 3 a), where 2’3’-cGAMP supplementation resulted in a clear, significant increase in autophagic flux, represented by clearly reduced p62 levels (Figure 5 c, d; Figure S5 b). In contrast, inhibition of STING by H-151 in DIV40 and DIV60 old neurons had no effect and accumulated in the soma comparable amounts of p62 relative to the untreated control group (Figure 5 c, d; Figure S5 b). Surprisingly, H-151 reduced the number of p62-positive puncta in the soma of DIV21 old neurons (Figure 5 d; Figure S5 a). The fact that additional CQ supplementation did not lead to an accumulation of p62 signals indicates a direct block of autophagy via the H-151-mediated inactivation of STING activity (Figure S5 c-d). Finally, we analyzed the influence of STING on lysosome biogenesis using the lysosome marker protein LAMP1. While activation of STING by 2’3’-cGAMP resulted in no effect in DIV21 old neurons (Figure 5 e; Figure S5 a), we observed a significant accumulation of LAMP1-positive lysosomes in the soma of DIV40 and DIV60 old neurons (Figure 5 c, e; Figure S5 b), indicating a direct involvement of STING in the lysosomal proteolytic pathway in aging neurons. When we blocked STING activity using H-151, we found a slight decrease in LAMP1 only in DIV21-old neurons, while its levels remained unchanged in DIV40 and DIV60-old neurons (Figure 5 c, e; Figure S5 a, b). From this we concluded that the activity of STING regarding the lysosome biogenesis in aging neurons is already restricted by the endogenous age-dependent reduced 2’3’-cGAMP levels.

**FIGURE 5.**
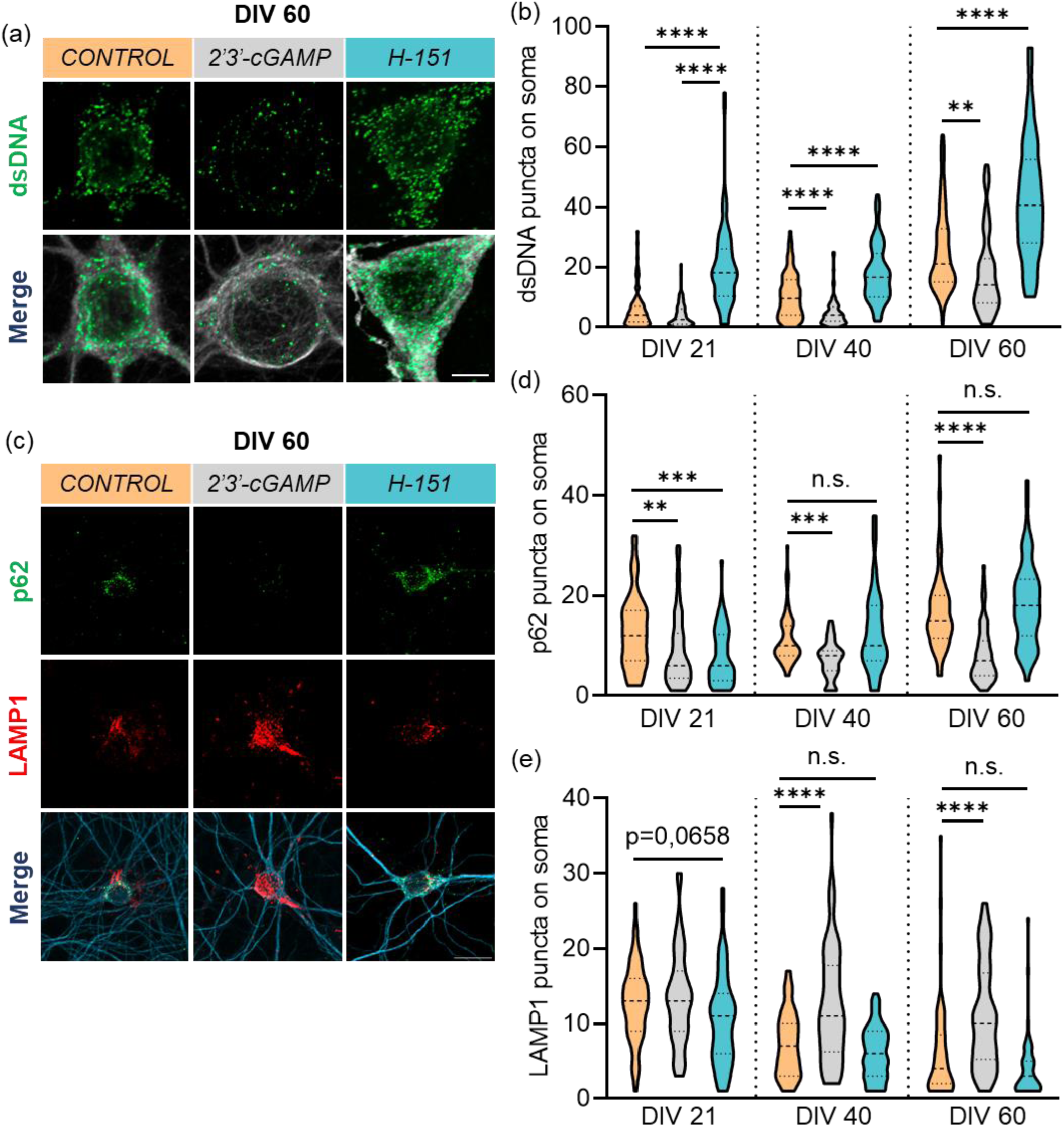
Induction of the cGAS-STING pathway reduces the age-dependent accumulation of cytosolic dsDNA and improves the autophagic status of old cortical neurons. (a) Representative images of dsDNA (green channel) on a MAP2 mask (white channel) without or with 2’3’-cGAMP (12,5 µg/mL) and H151 (4 µg/mL) treatment for 3 h on old cortical neurons DIV 60, scale bar = 5 µm. (b) Graph represents the number of dsDNA puncta evaluated on neuronal soma at DIV 21, 40 and 60 after activation or inhibition of STING activity. Graph shows n = 21-22 images/group from three independent experiments. (c) Immunofluorescence labelling of p62 (green channel), LAMP1 (red channel) and MAP2 (blue channel) without or with 2’3’-cGAMP (12,5 µg/mL) and H151 (4 µg/mL) treatment, scale bar = 20 µm. (d,e) Evaluation of p62 and LAMP1 puncta on neuronal soma at DIV 21, 40 and 60 without or with 2’3’-cGAMP and H151 treatment. Graph shows n = 19-25 images/group from three independent experiments. All data are presented as the mean ± SEM. Symbols for P-values used in the figures: \*\**P* < 0.01, \*\*\**P* < 0.001, \*\*\*\**P* < 0.0001. P-values were determined by ordinary one-way ANOVA with Tukey’s multiple comparisons.

### Age-related impaired phagosome-lysosome fusion in primary cortical neurons can be rescued by STING activation

Several reports suggest that the effectiveness and functionality of autophagic processes is directly related to the capacity of lysosomes to degrade the delivered cargo (for review see Carmona-Gutierrez et al., 2016). In addition, reduced loading and degradation in lysosomes appears to be involved in age-related processes and pathologies (Madeo et al., 2015, Sun et al., 2020). In fact, analysis of p62 and LAMP1 co-localization as a readout for autophagosome-lysosome fusion already showed a clear reduction in DIV40 old cultures and continued to decrease significantly up to DIV60 (Figure 5 c; 6 b; Figure S5 a, b), pointing towards a lysosomal dysfunction in aging neurons. Supplementation with 2’3’-cGAMP, the natural activator of STING, led to an almost complete restoration of autophagosome-lysosome fusion in DIV40 and DIV 60 neurons, while no effect could be observed in DIV21 neurons (Figure 6 a, c; Figure S5 a, b). On the one hand, this result indicates an age-dependent increasing impairment of autophagosome-lysosome fusion in neurons due to a simultaneously reduced STING signaling and, on the other hand, that young neurons at DIV21 have a sufficient STING signaling. These findings are supported by the observation that specific inhibition of STING activity by H-151 in young neurons at DIV21 resulted in a significant reduction of the autophagosome-lysosome fusion, while no changes could be shown in older neurons at DIV40 and DIV60 (Figure 6 a, c; Figure S5 a, b). Taken together, our results indicate a direct involvement of the cGAS-STING signaling pathway in autophagy-mediated lysosomal protein degradation in neurons and a significant impairment of these processes by an age-dependent loss of cGAS-STING activity.

**FIGURE 6.**
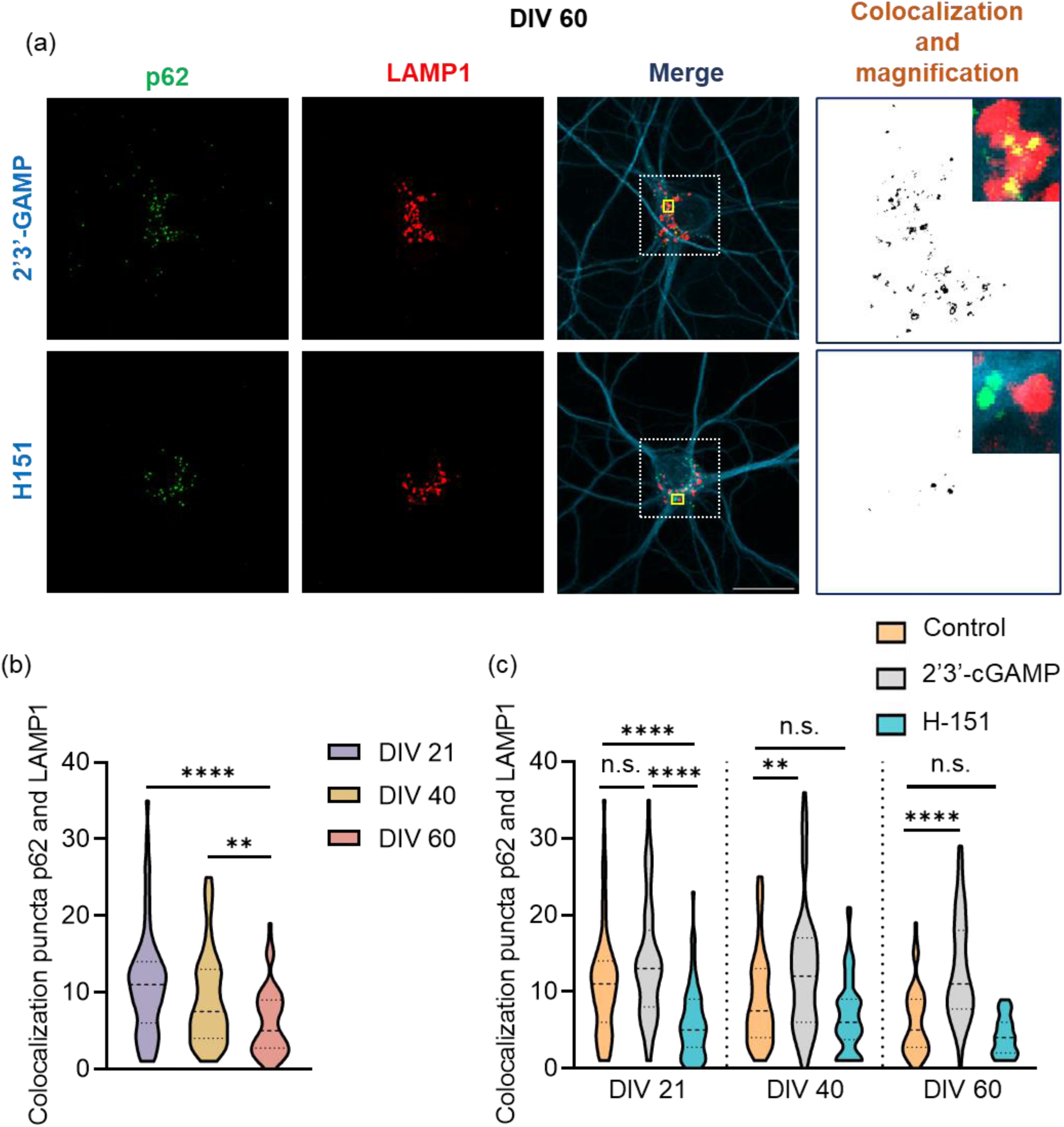
Rescue of the autophagosome-lysosome fusion after activation of STING activity in aged cortical neurons. (a) Representative figures of p62 (green channel), LAMP1 (red channel) and MAP2 (blue channel), scale bar = 20 µm. Colocalization analysis was performed with ImageJ’s Colocalization red and green tool. In addition, a magnification of an area of the neuronal soma is shown. (b) Quantification of p62 colocalization with LAMP1 on DIV 21, 40 and 60 neurons. Values are obtained from 17-18 figures done for each group from four independent experiments. (c) Quantification of p62 and LAMP1 colocalization without or with 2’3’-cGAMP and H151 treatment during aging *in vitro*. Graph shows n = 19-25 images/group from three independent experiments. All data are presented as the mean ± SEM. Symbols for P-values used in the figures: \*\**P* < 0.01, \*\*\*\**P* < 0.0001. P-values were determined by ordinary one-way ANOVA with Tukey’s multiple comparisons.

## 4. DISCUSSION

Healthy aging of the mammalian brain beyond age-related neurodegenerative diseases is still an elusive process demanding further intense research in this field. In our current study, we show that in neurons the role of cGAS-STING signaling is restricted to its primordial function of regulating autophagy and that this process is downregulated during neuronal aging. In contrast, we found no evidence in neurons that the cGAS-STING signaling pathway contributes significantly to IFN-I mediated inflammation, a process characteristic for microglia. Therefore, we hypothesize that the disability of neurons to generate noteworthy amounts of IFN-I during aging is part of its survival strategy as a postmitotic cell type.

To gain more insights into molecular changes of aging neurons, with particular emphasis on the regulation of autophagy, we developed an *in vitro* model system of primary neuronal cultures from mice cortices, consisting mainly of neurons and astrocytes, but lacking microglia, and led them age until DIV60. While in our cultures the concentrations of the dendritic and somatic marker MAP2 and the axonal marker TAU1 decreases continuously during the aging process *in vitro*, the steady increase of GFAP indicates both a sustained proliferation of the astrocytes and their increasing activation as it was demonstrated for the human cortex *in vivo* (Palmer & Ousman, 2018; Verkerke et al., 2021). In addition, we found in our cultures several senescence-and age-related markers that have been previously described both *in vivo* and *in vitro* (González-Gualda et al., 2021; Ishikawa & Ishikawa, 2020; Moreno-Blas et al., 2019; Sikora et al., 2014). For example, we observed a significantly increasing ß-Gal activity already at DIV40, indicative for senescence-related stress (Geng et al., 2010; Lee et al., 2006; Mera-Rodríguez et al., 2021), corresponding to increasing amounts of TUNEL positive and thus death cells, peaking at DIV60. Simultaneously, we detected an increasing accumulation of polyubiquitinated proteins, suggesting an age-dependent impairment of proteasomal protein degradation (Frankowska et al., 2022; Kelmer Sacramento et al., 2020). In addition, age-dependent increasing GST levels, with a simultaneous loss of mitochondria, most probably due to its malfunction and degeneration, points towards elevated concentrations of reactive oxygen species, as another marker of senescence (Fehrmann et al., 2013; Matoba et al., 2022). A more detailed cell type-specific analysis of the neurons in our cultures revealed a significant reduction in both dendritic branching and length after DIV60, consistent with reports from aging human and mammalian brains *in vivo* (D. L. Dickstein et al., 2013; Dara L. Dickstein et al., 2007; Hall et al., 2021; Müller-Thomsen et al., 2020; Tsai et al., 2018). Specifically, in neurons we also found significantly increased dsDNA levels as another hallmark of aging as early as DIV40, which nearly doubled by DIV60. Besides nuclear instability and uptake from the extracellular space via endocytosis, the mtDNA of the growing number of damaged mitochondria is supposed to be the main source for the dsDNA staining (Klein et al., 2021; Zia et al., 2021). In addition, we found an elevation of apoptotic processes in the neurons, which is evidenced by significantly increased levels of cleaved caspase 3, an indicator of final senescence processes (Su et al., 2001). In summary, we have succeeded in establishing an *in vitro* model of neuronal aging with up to 60-day-old neuronal primary cultures from the mouse cortex, which reflects essential elements of aging in the mammalian brain. This finding is also corroborated by the fact that we were able to show a significantly reduced protein synthesis in neurons using FUNCAT as early as DIV40, both in the soma and in the dendrites, which continued to decrease up to DIV60. Since impaired protein synthesis and degradation are other characteristics of senescence-related changes (Anisimova et al., 2018; Risi et al., 2020), we focused our further analysis on autophagy as an essential component of protein homeostasis. In our cultures we found a significant accumulation in both the lipidated form of LC3 (LC3/II) and an increase in the cargo receptor protein p62 up to DIV60. This result indicates a significant impairment of autophagic flux and is interpreted by several researchers as another feature of senescence (Ishikawa & Ishikawa, 2020; Moreno-Blas et al., 2019; Risi et al., 2020). A cell type-specific analysis of p62 in neurons also revealed a significant accumulation after DIV60, which indicates specifically in neurons an impairment of autophagic flux during aging. This finding is supported by the observation that after blocking autophagy using CQ, p62 accumulates only in DIV21 and DIV40 old neurons, but not in DIV60 old ones, pointing towards an already existing impairment of the autophagic flux in this aging group (Moreno-Blas et al., 2019). The age-dependently reduced autophagy in neurons is not only evident at the level of accumulating autophagosomes, but also by a significant decrease of LAMP1-positive lysosomes already after DIV40. The lysosomes at DIV60 appear significantly enlarged as demonstrated in previous research (Moreno-Blas et al. 2019). Since we also found an age-dependent elevation of lysotracker-positive lysosomes in neurons, we hypothesize that the proportion of functional lysosomes is increased during aging and accumulates because of prevented final processing. Furthermore, we simultaneously found in the aged neurons a reduction in the cis-Golgi complex, the cellular origin of lysosomes. In summary, we concluded an age-dependent impairment of the overall autophagy at different levels of the process.

Although autophagy in neurons, being postmitotic cells, is an extremely efficient mechanism with a rapid clearance of autophagosomes, little is known about its regulation in this cell type (Yang & Klionsky, 2020). In fact, in recent years there is an increasing amount of research demonstrating autophagy impairment in neurodegenerative disease such as Alzheimer’s disease (AD), Parkinson’s disease (PD) or Huntington’s disease (HD), and during healthy aging (Choi et al., 2020; Heckmann et al., 2020; Lipinski et al., 2010; Lizama & Chu, 2021; Oh et al., 2022; Reddy & Oliver, 2019). Further elucidation of the role of autophagy and its regulation in neurons will be key to a better understanding of age-related neuronal dysfunction and degeneration (Magalhães et al., 2021). Therefore, since we observed a significantly increasing accumulation of dsDNA in the soma of aging neurons, we asked the question to what extent noncanonical, STING-dependent autophagy plays a role in the age-related impairment of neuronal autophagy. In this context, we were able to show that in our cultures the amount of the dsDNA sensor cGAS dropped significantly to around 20% of the initial concentration after DIV60, while the amount of STING remained almost constant over the same period. The decline of cGAS can be explained by the age-dependent increase in cleaved caspase 3 in neurons that we observed. Indeed, active apoptotic caspases, like caspase 3, can negatively modulate the cGAS-STING pathway due to cleaving of cGAS protein (Ning et al., 2019). As a direct consequence, the 2’3’-cGAMP concentration in our cultures also decreased as a function of age. Blocking of STING activity using H-151 showed that the ability of cGAS to produce 2’3’-cGAMP was significantly restricted at DIV60, and the activity of the cGAS-STING signaling pathway was thus down-regulated overall. In this context, we detected a slight reduction of the pTBK1 concentration and a significant decrease of pIRF3 to approximately 25% of the initial level at DIV60, which is consistent with the observed decrease in cGAS concentration and activity. Since increased levels of pIRF3 are a basic requirement for interferon type I expression induced via the cGAS-STING signaling pathway (Sun et al., 2013), which is subsequently responsible for activating mechanisms of innate immunity (X.-D. Li et al., 2013), we hypothesize that specifically this function is not primary or negligible for the importance of the cGAS-STING signaling pathway in aging neurons. This assumption is consistent with previous reports showing that the activity of caspases prevents mtDNA-dependent IFN-I production (Rongvaux et al., 2014; White et al., 2014).

To gain more insights into the functioning of STING within the regulation of the non-canonical autophagy machinery in neurons, we analyzed first its expression in neurons and were able to confirm this by using specific antibodies and primary neurons from wildtype mice and STING-KO mice as a control, what is in line with a couple of other publications (Abdullah et al., 2018; Donnelly et al., 2021). The activation of STING using its endogenous agonist 2’3’-cGAMP resulted in a significant reduction of the dsDNA levels in the soma of both DIV40 and DIV60 old neurons, while no effect was observed in young neurons. This suggests that in young neurons, adequate levels of 2’3’-cGAMP can mediate sufficient clearance via the cGAS-STING axis, while decreasing 2’3’- cGAMP levels during aging inactivate this pathway. Blocking of STING activity using the specific inhibitor H-151 led to a significant accumulation of dsDNA in the soma of the neurons in all age groups examined and proves the direct involvement of cGAS-STING mediated autophagy in processes of neuronal proteostasis. This finding is supported by a direct analysis of autophagosomes in the soma of aging neurons, whose accumulation was almost eliminated after activation of STING, while the amount of autophagosomes accumulated after inhibition of STING remained at the level of the control group. The fact that inactivation of STING in young neurons leads to lower levels of p62 indicates a direct influence of STING on autophagosome biogenesis, at least at this stage of development.

The increasing number of LAMP1-positive lysosomes and p62-LAMP1 fusions in aging neurons after activation with 2’3’-cGAMP showed that STING also has an impact on both lysosomal biogenesis and the autophagosome-lysosome fusion rate. Both the number of LAMP1-positive lysosomes and the level of autophagosome-lysosome fusions in DIV40 and DIV60 old neurons reached the values observed in young neurons and thus corresponds to an almost complete rescue, which proves a direct dependency on the activity of the cGAS-STING axis. Inhibition of STING activity in young neurons prevented autophagosome-lysosome fusion but, in contrast, had no effect on DIV40 and DIV60 old neurons, suggesting an endogenously age-dependent downregulated activity of the cGAS-STING axis. Summarizing the latter points, we hypothesize that cGAS-STING signaling is significantly involved in the final steps of autophagy, lysosome biogenesis and autophagosome-lysosome fusion, and thus makes an essential contribution to neuronal proteostasis. This finding goes in line with previous reports, demonstrating autophagy induction via IFN-independent STING signaling (Gui et al, 2019, Yamashiro et al, 2020, Wan et al. 2023).

In summary, we present an *in vitro* model to study the increasingly impaired proteostasis in aging neurons and showed for the first time that individual stages of autophagy-dependent cellular clearance in neurons are affected by decreasing cGAS-STING signaling activity. Thereby, STING-mediated signaling appears to affect both the initial biogenesis of autophagosomes, probably due to its WIPI2 interaction (Gui et al, 2019, Wan et al. 2023), and the biogenesis and fusion of lysosomes. In addition, we could show significantly decreasing levels of pIRF3 in the aged cells, which can be attributed to a simultaneous accumulation of activated caspase 3 in neurons. This points to the fact that in contrast to microglia, aging neurons do not contribute to an IFN-I-dependent inflammatory process, despite constantly accumulating amounts of dsDNA as an activator of the cGAS-STING signaling pathway. Considering these data, we hypothesize that in neurons the cGAS-STING signaling pathway is primarily involved in the regulation of noncanonical autophagy processes contributing to the cellular homeostasis and seems to be not engaged as a mediator of inflammation and innate immunity. Potentially, this represents a specific higher-order survival strategy of postmitotic neurons and thus distinguishes them from other cells of the brain, e.g., microglia.

## Supporting information

Supporting informations

## ACKNOWLEDGMENTS

This work was supported by SFB 854, RTG 2413 Synage and Research Unit 5228 Syntophagy-DI1512/7-1.

We thank Saskia Kresse, I-Chin Tsai and Parthiban Saravanakumar for the technical assistance, Dr. Anke Müller for the fruitful discussions for the execution of certain experiments, Kathrin Freke for animal care taking, Prof. Dr. Anne Albrecht and Dr. Stefan Kahlert for the support in imaging.

## AUTHOR CONTRIBUTION

SP was involved in conception and study design, conducted experiments, analyzed, interpreted and discussed data, manuscript writing. KB, ED, PC were involved in conducted experiments and maintenance of cultures. AK, DCD were involved in conception and study design. DCD supervised the study and was also involved in manuscript writing, discussed data and provided financial support. PL conceptualized and supervised the study, interpreted, discussed data and wrote the manuscript.

## DATA AVAILABILITY STATEMENT

The data are available from the corresponding author upon reasonable request.

