## Supporting informations for "Neuronal aging is associated with declined autophagy caused by reduced functionality of cGAS-STING signaling"

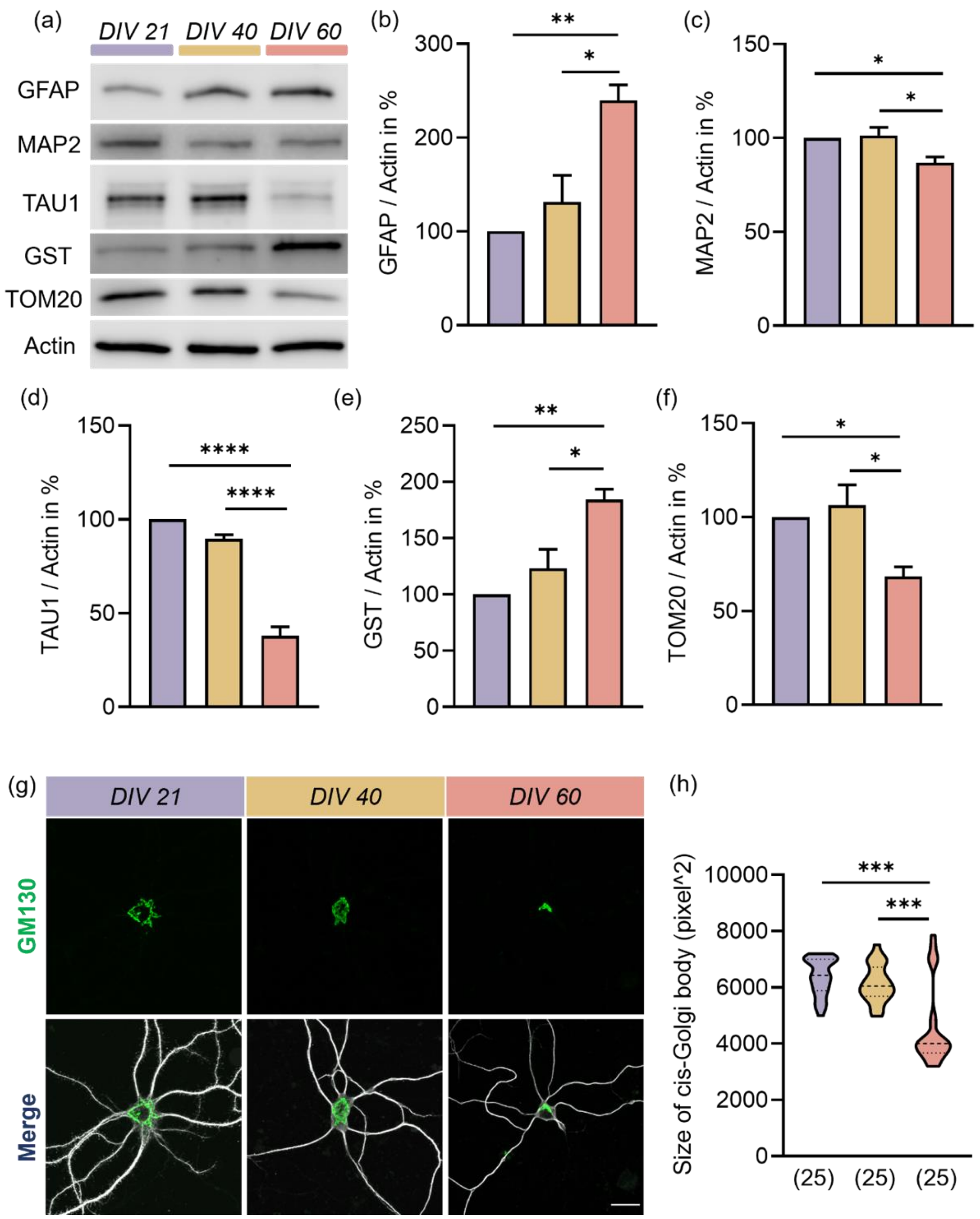


**Supplementary figure 1**: Quantitative analysis of neural and cellular marker in primary cortical cultures at different ages (a) In order to characterize the cell population and the cell stress in culture immunoblots of selected GFAP, MAP2, TAU1 and GST protein were performed. (b-f) Relative quantities of Western blot analysis of GFAP, MAP2, TAU1, GST and TOM20 normalized with Actin. Values are the mean of three technical replicates in each group from three independent primary neural cultures for immunoblotting analysis. (g,h) Neurons were stained with GM130 (green channel) and MAP2 (white channel) at different time point DIV 21, 40 and 60 and evaluation of the Golgi size measured in pixel$^2$, scale bar = 20 µm. Graph n = 8 - 9 images done for each group from three independent experiments for immunostainings analysis. All data are presented as the mean ± SEM. Symbols for P-values used in the figures: *$P < 0.01$, **$P < 0.01$, ***$P < 0.001$, ****$P < 0.0001$. P-values were determined by ordinary one-way ANOVA with Tukey's multiple comparisons test.

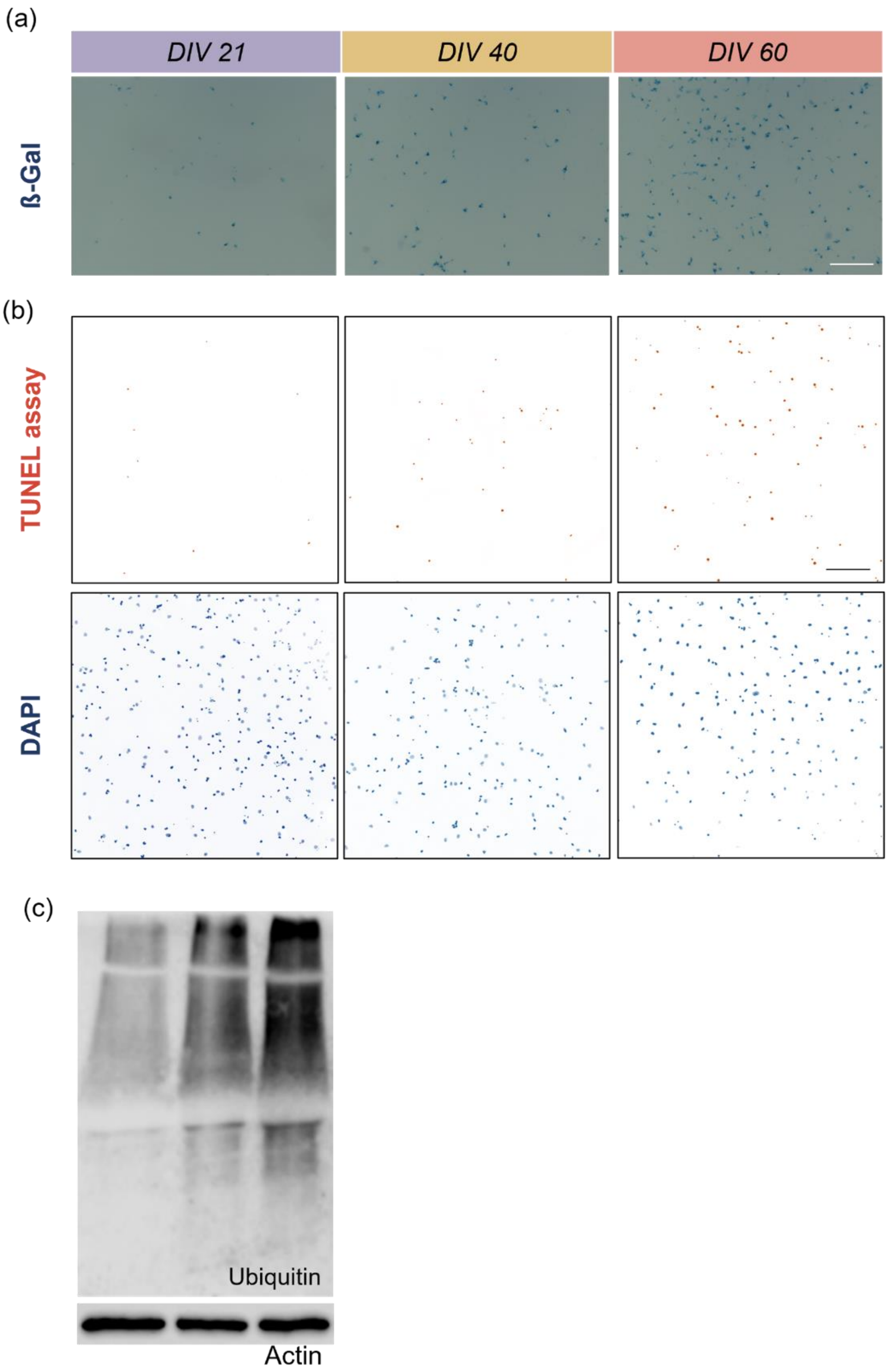


**Supplementary figure 2:** (a) Primary neural mice cultures were stained with beta-galactosidase in order to evaluate a senescent feature at different time points (DIV 21, DIV 40 and DIV 60) on pictures with a size of 2195 µm x 1737 µm, scale bar: 100 µm. (b) Cells were stained with TUNEL assay (FITC, red) for the quantification of the fragmented DNA. The signal of DAPI is marked in blue, scale bar: 50 µm. The size of TUNEL assay pictures is 850,19 x 850,19 µm. (c) Representative blots of Ubiquitin and the control protein Actin. These results are based on 3 independent experiments, run in triplicates

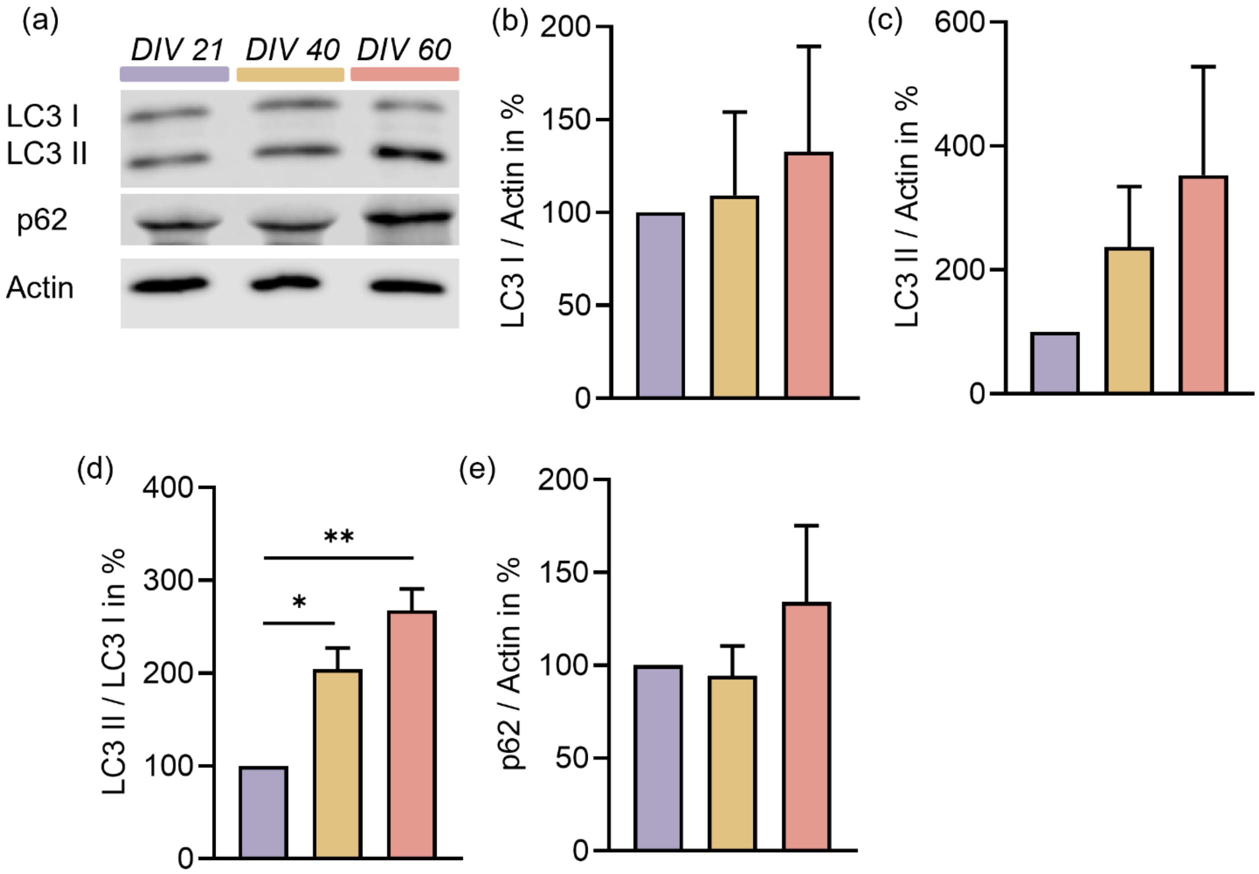


**Supplementary figure 3:** Aging of neural cells *in vitro* leads to an impairment of the autophagic flux. (a) Immunoblotting of LC3 and p62 with Actin used as control from three independent cultures of neural cells at DIV 21, 40 and 60. (b-e) Quantitative data of LC3 I, LC3 II, ratio LC3 II / LC3 I and p62 normalized to Actin. Values are the mean of three technical replicate in each group from three independent experiments for immunoblotting analysis. All data are presented as the mean ± SEM. Symbols for P-values used in the figures: $^{*}P < 0.05$, $^{**}P < 0.01$, $^{***}P < 0.001$, $^{****}P < 0.0001$. P-values were determined by ordinary one-way ANOVA with Tukey's multiple comparisons test.

(a)

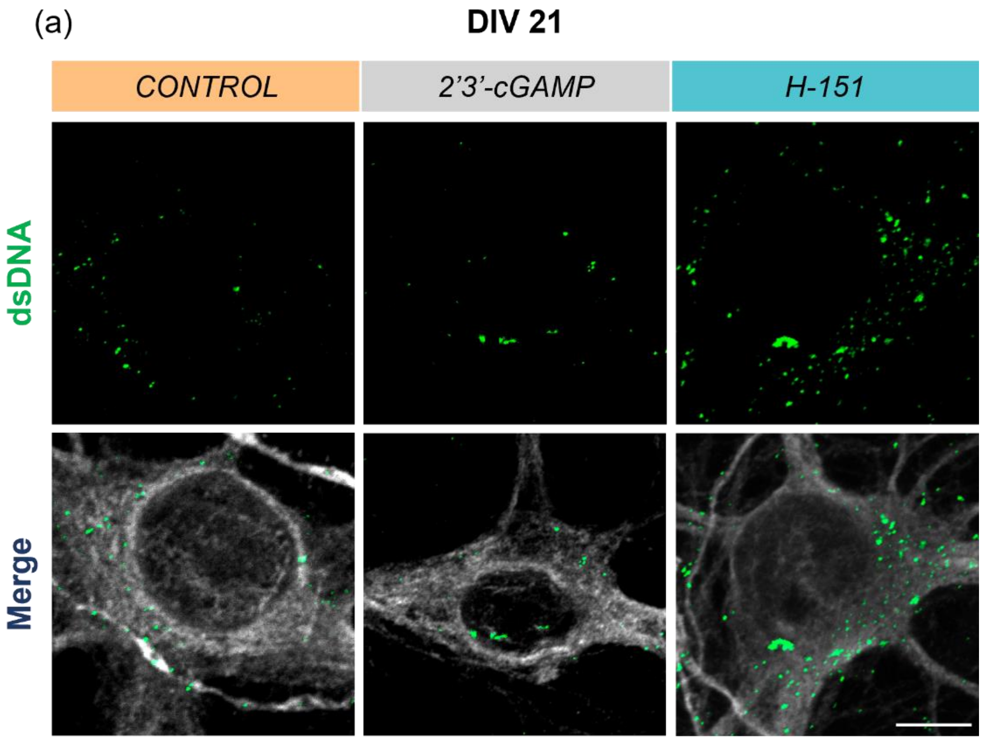


(b)

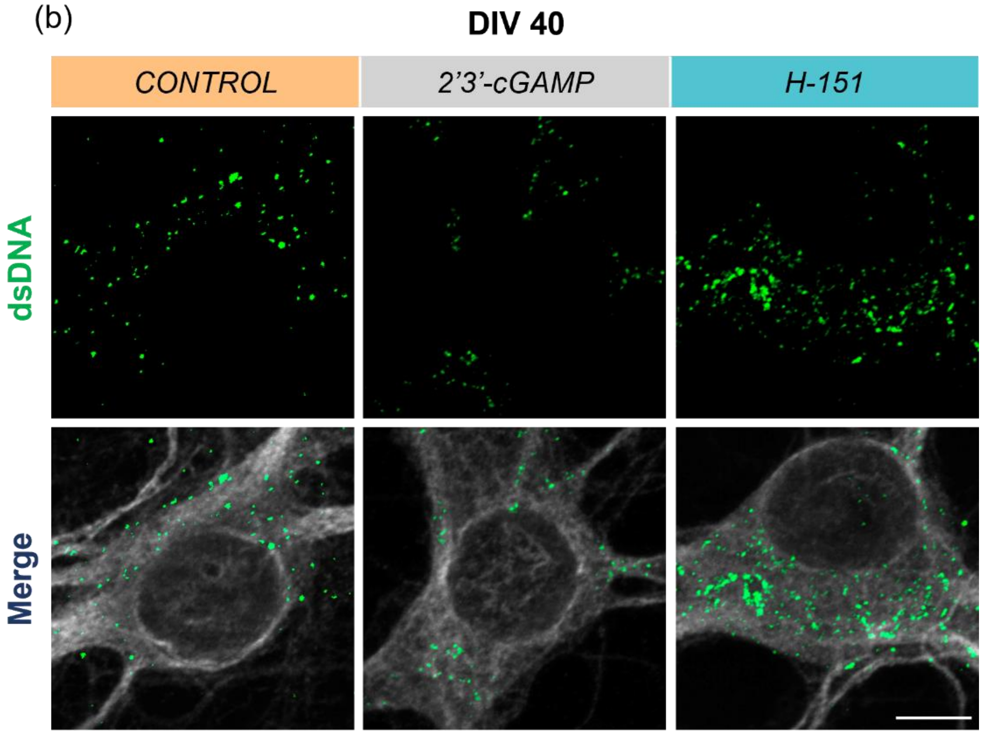


**Supplementary figure 4:** (a,b) Representative pictures of dsDNA (green channel) staining on mouse cortical neurons labelled with MAP2 (white channel) at DIV 21 and 40, scale bar = 20 µm. The groups analyzed were control, 2'3'-cGAMP (12,5 µg/mL) and H-151 (4 µg/mL).

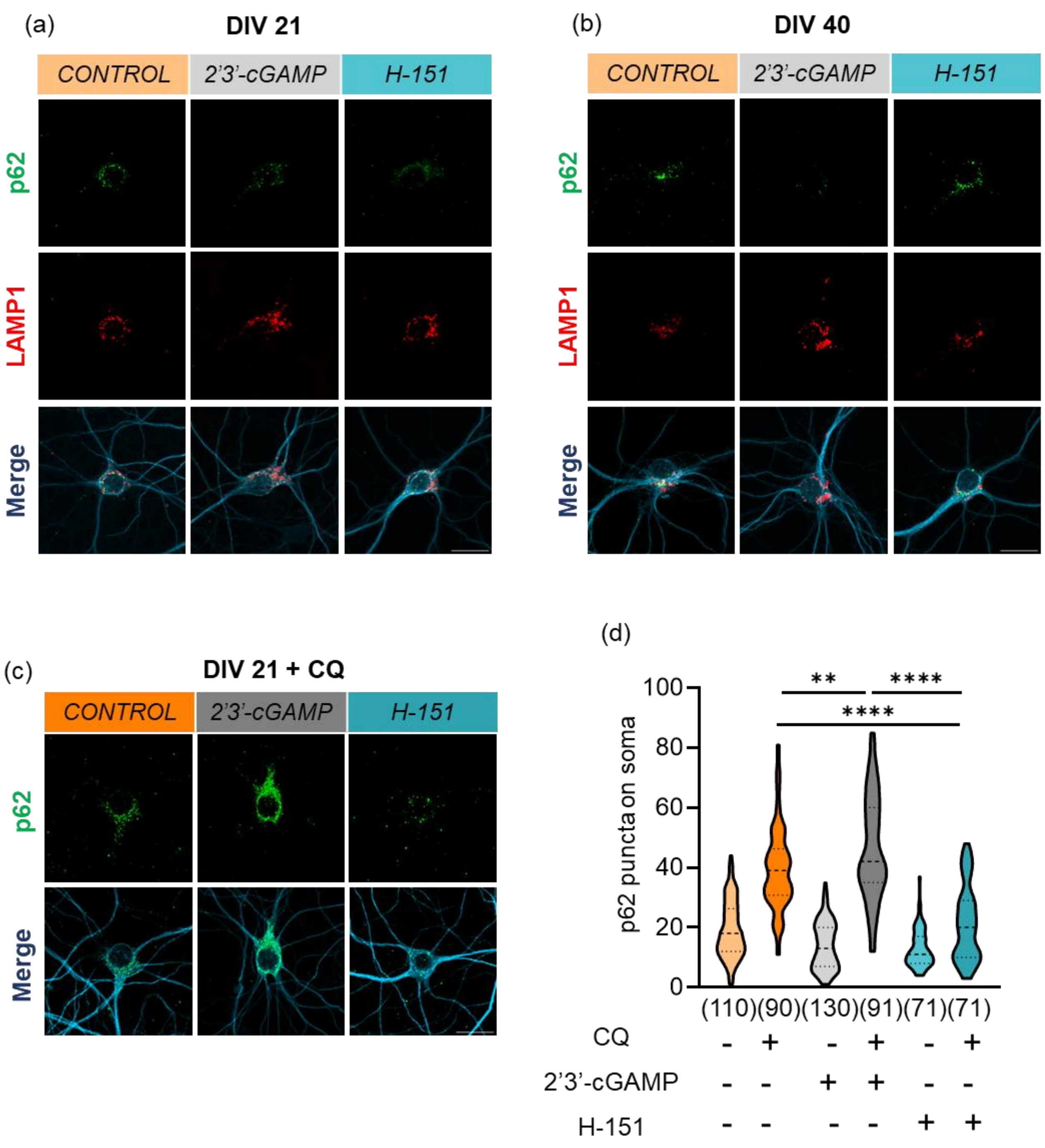


**Supplementary figure 5:** (a,b) Immunofluorescence labelling of p62 (green channel), LAMP1 (red channel) and MAP2 (blue channel) without or with 2'3'-cGAMP (12,5 µg/mL) and H151 (4 µg/mL) treatment of neurons at DIV 21 and 40, scale bar = 20 µm. (c-d) Quantification of p62 puncta on mature neurons at DIV 21. The groups analyzed were control, 2'3'-cGAMP (12,5 µg/mL) and H-151 (4 µg/mL) with or without a co-treatment with 50 µM CQ. Graph shows n = 25-40 images/group from three independent experiments. All data are presented as the mean ± SEM. Symbols for P-values used in the figures: *$P < 0.05$, **$P < 0.01$, ***$P < 0.001$, ****$P < 0.0001$. P-values were determined by ordinary one-way ANOVA with Tukey's multiple comparisons test.
